# Polymerization of C9 enhances bacterial cell envelope damage and killing by membrane attack complex pores

**DOI:** 10.1101/2021.05.12.443779

**Authors:** Dennis J. Doorduijn, Dani A.C. Heesterbeek, Maartje Ruyken, Carla J.C. de Haas, Daphne A.C. Stapels, Piet C. Aerts, Suzan H. M. Rooijakkers, Bart W. Bardoel

## Abstract

Complement proteins can form Membrane Attack Complex (MAC) pores that directly kill Gram-negative bacteria. MAC pores assemble by stepwise binding of C5b, C6, C7, C8 and finally C9, which can polymerize into a transmembrane ring of up to 18 C9 monomers. It is still unclear if the assembly of a polymeric-C9 ring is necessary to sufficiently damage the bacterial cell envelope to kill bacteria, because a robust way to specifically prevent polymerization of C9 has been lacking. In this paper, polymerization of C9 was prevented without affecting the binding of C9 to C5b-8 by locking the first transmembrane helix domain of C9. We show that polymerization of C9 strongly enhanced bacterial cell envelope damage and killing by MAC pores for several *Escherichia coli* and *Klebsiella* strains. Moreover, we show that polymerization of C9 is impaired on complement-resistant *E. coli* strains that survive killing by MAC pores. Altogether, these insights are important to understand how MAC pores kill bacteria and how bacterial pathogens can resist MAC-dependent killing.

## Introduction

Complement proteins in human serum play a crucial role in fighting off invading bacteria. Activation of the complement cascade ultimately results in the assembly of membrane attack complex (MAC) pores that can directly kill Gram-negative bacteria [1–3]. MAC assembly is initiated when recognition molecules, such as antibodies and lectins, bind to bacteria and recruit early complement proteins [4]. This triggers a proteolytic cascade that deposits convertase enzymes on the bacterial surface [5]. These convertases convert complement component C5 into C5b, which initiates the assembly of a large ring-shaped MAC pore that damages the bacterial cell envelope [3,6,7]. Although MAC pores can efficiently kill complement-sensitive bacteria, some bacterial pathogens can survive killing by MAC pores [8–11]. Therefore, studying how MAC pores kill bacteria is important to understand how the complement system prevents infections and how bacterial pathogens resist killing by MAC pores.

MAC pores assemble in a stepwise manner [12]. When a surface-bound convertase converts C5 into C5b, C5b immediately binds to C6 to form the C5b6 complex [13,14]. Direct binding of C7 to C5b6 stably anchors the MAC precursor to the membrane [15]. Next, C8 binds to membrane-anchored C5b-7, which triggers structural rearrangements in C8 that result in insertion of a transmembrane β-hairpin into the membrane [16,17]. Finally, C9 binds to C5b-8 and polymerizes to form a transmembrane ring of up to 18 monomers C9 with an inner diameter of 11 nm [18,19].

Although C5b-8 can already cause small 1-2 nm lesions in the membrane of erythrocytes and liposomes without a polymeric-C9 ring [17,20], it is still unclear if C9 polymerization is required to sufficiently damage the complex bacterial cell envelope to kill Gram-negative bacteria. On bacteria, MAC pores initially assemble on the outer membrane (OM), which largely consists of lipopolysaccharide (LPS). The O-antigen (O-Ag) of LPS can vary in length between bacterial strains and species [21,22], and this has frequently been associated with complement-resistance [8,9,23,24]. Apart from the OM, the Gram-negative cell envelope also consists of a cytosolic inner membrane (IM) and a periplasmic peptidoglycan layer [25]. We recently developed methods to separately study OM and IM damage in time [14,26]. Here, we wanted to use these methods to study how C9 polymerization contributes to bacterial cell envelope damage by MAC pores. Moreover, a direct causal link between polymerization of C9 and bacterial killing has still not been established, mainly because there was no robust system in which polymerization of C9 could specifically be prevented. Recently, Spicer *et al*. suggested that ‘locking’ the first transmembrane helix (TMH-1) domain of C9 could prevent polymerization of C9, without affecting the binding of C9 to C5b-8 [27]. Here, we wanted to use this ‘locked’ C9 to study if C9 polymerization contributes to bacterial cell envelope damage and killing by MAC pores.

In this paper, we show that polymerization of C9 enhanced the efficiency by which MAC pores damage both the OM and IM, which ultimately resulted in faster killing of several *Escherichia coli* and *Klebsiella* strains. This study therefore highlights that MAC pores have to completely assemble to efficiently damage the bacterial cell envelope and kill bacteria. Moreover, we found that polymerization of C9, but not binding of C9 to C5b-8, was impaired on several complement-resistant *E. coli* strains that survive killing by MAC pores. This study therefore also provides insights into how bacterial pathogens resist MAC-dependent killing.

## Results

### Locking the TMH-1 of C9 strongly impairs its capacity to polymerize, without preventing binding to C5b-8 on *E. coli*

To study the contribution of C9 polymerization to bacterial killing by the MAC, we wanted to use a system in which C9 can bind to C5b-8, but cannot polymerize. Spicer *et al.* recently suggested that the TMH-1 domain of C9 has a crucial role in C9 polymerization [27]. When C9 binds to C5b-8, structural rearrangements in C9 trigger unfurling of the TMH-1 (**Fig. 1a-I**) and TMH-2 (not shown in illustration) to form a transmembrane β-hairpin. Although both TMH domains of C9 insert into the membrane, unfurling of the TMH-1 domain also exposes an elongation surface that allows a subsequent C9 to bind (**Fig. 1a-II**). This ultimately results in the assembly of a polymeric-C9 ring (**Fig. 1a-III)**. Based on this crucial role of the TMH-1 domain in polymerization, Spicer *et al.* designed a C9 TMH-1 ‘lock’ mutant (C9_TMH-1 lock_) in which the TMH-1 domain was linked to β-strand 4 of the MACPF/CDC domain via an intramolecular cysteine bridge. This lock prevents unfurling of the TMH-1 domain once C9 binds to C5b-8, and thus prevents both the formation of a transmembrane β-hairpin (**Fig. 1a-IV**) and binding of a subsequent C9 (**Fig. 1a-V**). Reducing the cysteine bridge with DTT can unlock the TMH-1 domain and restore its capacity to polymerize (**Fig. 1a-VI**).

**Figure 1.**
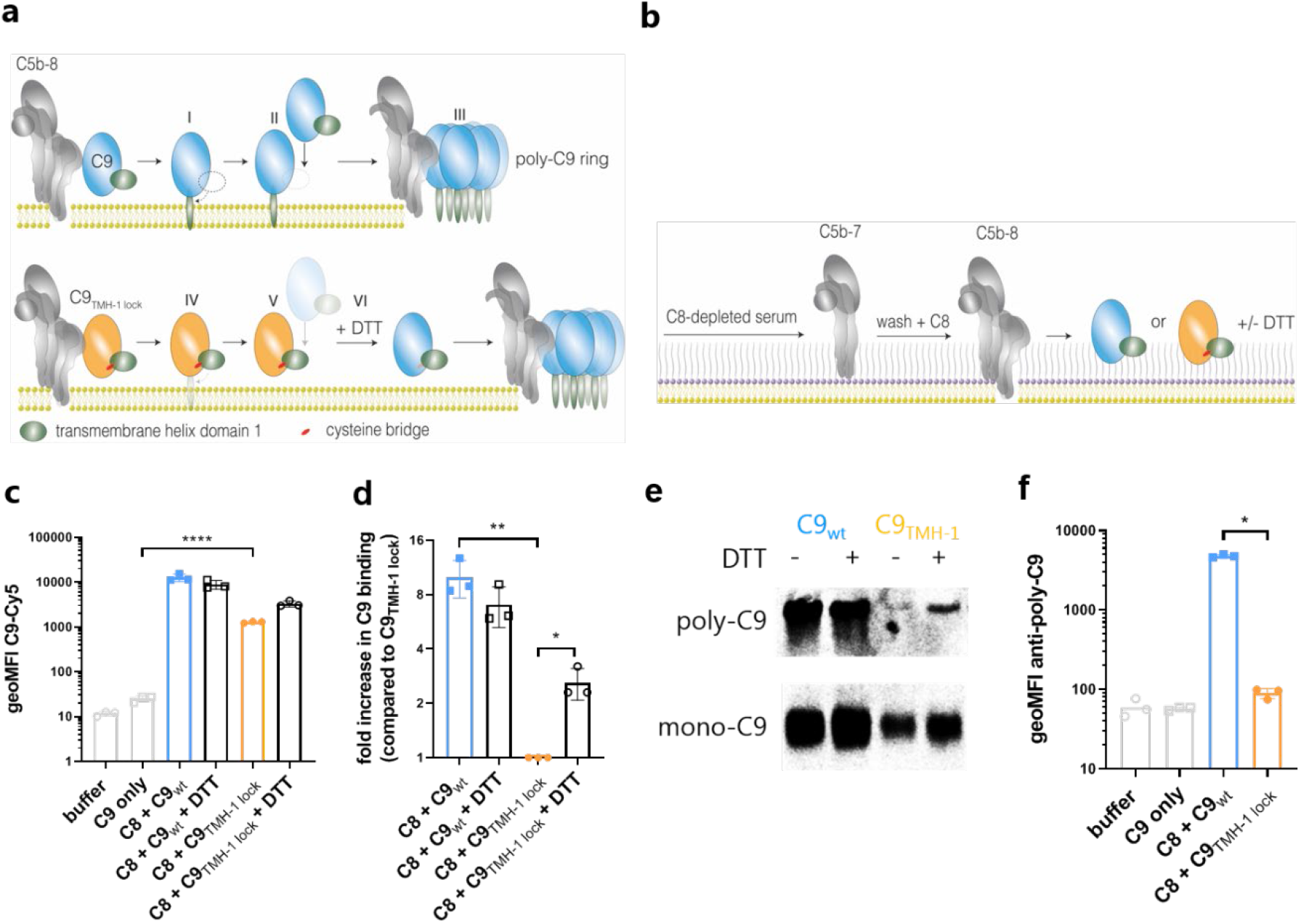
C9_TMH-1 lock_ binds to C5b-8, but its capacity to form polymers is impaired on *E. coli*. a) Schematic overview of C9 polymerization. Binding of C9 (blue) to C5b-8 (grey) triggers unfurling of the TMH-1 domain of C9 (green) and subsequent insertion into the membrane (I). Unfurling of TMH-1 exposes the elongation surface of C9 to bind a subsequent C9 monomer (II). Ultimately, this results in the formation of a C9 polymer (III). Locking the TMH-1 domain of C9 (orange, C9_TMH-1 lock_) with an intramolecular cysteine bridge (red) prevents unfurling of the TMH-1 domain when C9 binds to C5b-8 (IV). This also prevents exposure of the elongation surface of C9_TMH-1 lock_ and subsequent polymerization (V). Reducing the cysteine bridge of C9_TMH-1 lock_ with DTT restores its capacity to form polymers (VI). b) Schematic overview of labelling *E. coli* MG1655 with MAC. *E. coli* is labelled with C5b-7 by incubating them in 10% C8-depleted serum for 30 minutes. Bacteria are washed and next incubated with 10 nM C8 for 15 minutes. Finally, 20 nM of Cy5-labelled C9_wt_ or C9_TMH-1 lock_ is added in the presence or absence of 10 mM DTT for 30 minutes to measure C9 binding. c) Binding of Cy5-labelled C9_wt_ or C9_TMH-1 lock_ to bacteria treated as described in b and measured by flow cytometry. d) Binding of Cy5-labelled C9 in c was divided by the fluorescence of bacteria labelled with C8 + C9_TMH-1 lock_ to calculate the relative binding difference as indication for polymerization of C9. e) Bacterial cell pellets were analyzed by SDS-PAGE for in-gel fluorescence of Cy5-labelled C9_wt_ or C9_TMH-1 lock_ to distinguish monomeric-C9 (mono-C9) from polymeric-C9 (poly-C9). f) Bacteria were stained with AF488-labelled mouse anti-poly-C9 aE11-antibody and staining was measured by flow cytometry. Flow cytometry data are represented by individual geoMFI values of the bacterial population. SDS-PAGE images are representative for at least three independent experiments. Data represent individual values of three independent experiments with mean +/− SD. Statistical analysis was done on log-transformed data (^10^log for c, f and ^2^log for d) using a paired one-way ANOVA with Tukey’s multiple comparisons’ test. Significance was shown as * p ≤ 0.05, ** p ≤ 0.005, **** p ≤ 0.0001.

C9_TMH-1 lock_ was recombinantly expressed and site-specifically labelled with a fluorophore via sortagging, as was done previously for wildtype C9 (C9_wt_) [14]. Fluorescent labelling was comparable between both C9_TMH-1 lock_ and C9_wt_ (**S1a**), which means that the fluorescence of both proteins can be directly compared in our assays. C9_TMH-1 lock_ showed impaired lysis of sheep erythrocytes compared to C9_wt_, which could be restored by reducing C9_TMH-1 lock_ with DTT (**S1b**). Moreover, fluorescent C9_TMH-1 lock_ was used to distinguish SDS-stable polymeric-C9 (poly-C9) from monomeric-C9 (mono-C9) by SDS-PAGE, which is frequently used as a read-out for C9 polymerization [28]. C9_TMH-1 lock_ did not form poly-C9 together with preassembled C5b6 (pC5b6), C7 and C8, whereas C9_wt_ did (**S1c**). These data confirm that the capacity of C9_TMH-1 lock_ to form polymers is impaired, and can be reversed by reducing the cysteine bridge lock.

Next, we wanted to validate that locking the TMH-1 domain only prevents polymerization of C9, without preventing binding of C9 to C5b-8. *E. coli* MG1655 bacteria were incubated in C8-depleted serum to activate complement and label them with MAC precursor C5b-7 (**Fig. 1b**). C5b-7 labelled bacteria were washed to remove remaining serum components and incubated with C8 and C9_wt_ or C9_TMH-1 lock_ to further assemble the MAC. Both C9_wt_ and C9_TMH-1 lock_ bound to C5b-7 in a C8-dependent manner as measured by flow cytometry (**Fig. 1c**), although C9_wt_ binding was 10-fold higher than C9_TMH-1 lock_. Since the amount of C5b-8 on the surface, as measured by C6-FITC binding, was comparable for both C9_wt_ and C9_TMH-1 lock_ (**S1d**), the relative difference in C9 binding suggested a difference in polymerization of C9 (**Fig. 1d**). SDS-PAGE confirmed that only mono-C9 was detected on bacteria incubated with C9_TMH-1 lock_ (**Fig. 1e**). Moreover, reducing C9_TMH-1 lock_ with DTT increased binding 3-fold compared to C9_TMH-1 lock_ without DTT (**Fig. 1c,d**) and resulted in the detection of poly-C9 on bacteria by SDS-PAGE (**Fig. 1e**). This suggests that reducing C9_TMH-1 lock_ only partially restored its capacity to polymerize compared to C9_wt_ (**Fig. 1c,d**). Finally, an antibody that recognizes a neo-epitope exposed in poly-C9, which is frequently used for the detection of MAC pores [29], specifically detected bacteria incubated with C9_wt_, but not with C9_TMH-1 lock_ (**Fig. 1f**). Altogether, these data indicate that the C9_TMH-1 lock_ can bind to C5b-8 on *E. coli*, but that its capacity to polymerize is strongly impaired.

### Polymerization of C9 enhances bacterial killing by MAC pores

We then assessed if polymerization of C9 is important for bacterial killing by the MAC. A DNA dye that cannot permeate an intact IM (Sytox) was used to measure the percentage of cells with IM damage by flow cytometry, which we have previously shown to be a sensitive read-out for bacterial killing [14]. Adding C9_wt_ to C5b-8 labelled bacteria resulted in IM damage in a dose-dependent manner, reaching 100% Sytox positive cells from above 3 nM C9 (**Fig. 2a**). For C9_TMH-1 lock_, IM damage was impaired and did not increase above 30% Sytox positive cells at 100 nM C9 (**Fig. 2a**). Moreover, bacterial viability was determined by counting colony forming units (CFU’s) and decreased only 10-fold for C9_TMH-1 lock_ compared to C5b-8 alone, whereas C9_wt_ decreased bacterial viability at least a 1,000-fold (**Fig. 2c**). Reducing C9_TMH-1 lock_ with DTT restored its capacity to damage the IM (**Fig. 2b**) and kill bacteria (**Fig. 2c**). Finally, polymerization of C9_wt_ (**Fig. 2d**) and subsequent IM damage (**Fig. 2e**) could be inhibited by C9_TMH-1 lock_ in a dose-dependent manner. Poly-C9 detection already decreased by 50% when the amount of C9_TMH-1 lock_ was still 10-fold lower than the amount of C9_wt_ (**Fig. 2d**), which suggested that C9_TMH-1 lock_ can interfere at multiple stages in the assembly of a polymeric-C9 ring. However, IM damage was only fully inhibited when there was 10-fold more C9_TMH-1 lock_ than C9_wt_ (**Fig. 2e**), suggesting that only very few C9 polymers are required to damage the IM. Altogether, our data suggest that polymerization of C9 enhances bacterial killing by MAC pores.

**Figure 2.**
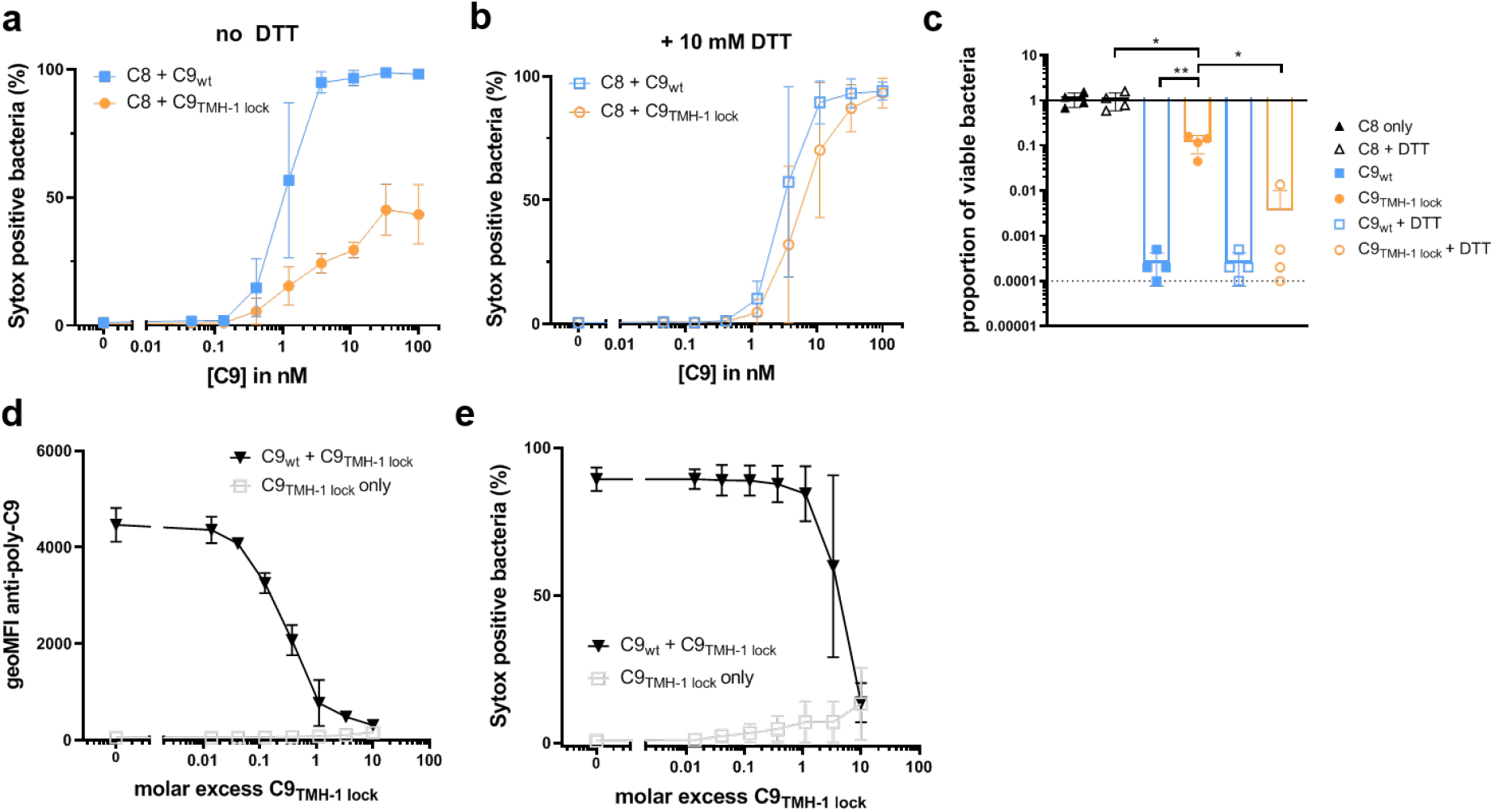
Polymerization of C9 enhances bacterial killing by MAC pores. *E. coli* MG1655 were labelled with C5b-7 by incubating them in 10% C8-depleted serum for 30 minutes. Bacteria were washed and next incubated with 10 nM C8 for 15 minutes. a-c) A concentration range of C9_wt_ or C9_TMH-1 lock_ was added in the absence (a) or presence (b) of 10 mM DTT for 30 minutes. Sytox was used to determine the percentage of cells that have a damaged bacterial IM by flow cytometry as read-out for bacterial killing. c) At 100 nM C9, bacterial viability was determined by counting colony forming units (CFU’s) and calculating the proportion of viable cells compared to C5b-7 labelled bacteria in buffer. The horizontal dotted line represents the detection limit of the assay. d-e) C5b-8 labelled bacteria were incubated for 30 minutes with 20 nM C9_wt_ and a concentration range of C9_TMH-1 lock_. Bacteria were stained with AF488-labelled mouse anti-poly-C9 aE11-antibody (d) and Sytox to determine the percentage of cells that has a damaged bacterial IM (e) by flow cytometry. Flow cytometry data are represented by geoMFI values or cell frequencies of the bacterial population. Data represent mean values +/− SD (a,b,d,e) or individual values (c) of three independent experiments with mean +/− SD. Statistical analysis was done on ^10^log-transformed data (c) using a paired one-way ANOVA with Tukey’s multiple comparisons’ test. Significance was shown as * p ≤ 0.05, ** p ≤ 0.005.

### Polymerization of C9 increases OM damage

Since MAC pores initially assemble on the OM [14], we also wondered how C9 polymerization contributes to OM damage. First, OM damage was compared for C9_wt_ and C9_TMH-1 lock_ by measuring leakage of periplasmic mCherry (_peri_mCherry, 22 kDa) through the OM of MG1655 using flow cytometry [14]. Adding C9_wt_ to C5b-8 labelled bacteria resulted in rapid leakage of _peri_mCherry within 5 minutes (**Fig. 3a**). By contrast, no more _peri_mCherry leaked out with C9_TMH-1 lock_ compared to C5b-8 alone within 60 minutes, suggesting that polymerization of C9 is required to cause leakage of periplasmic proteins through the OM.

**Figure 3.**
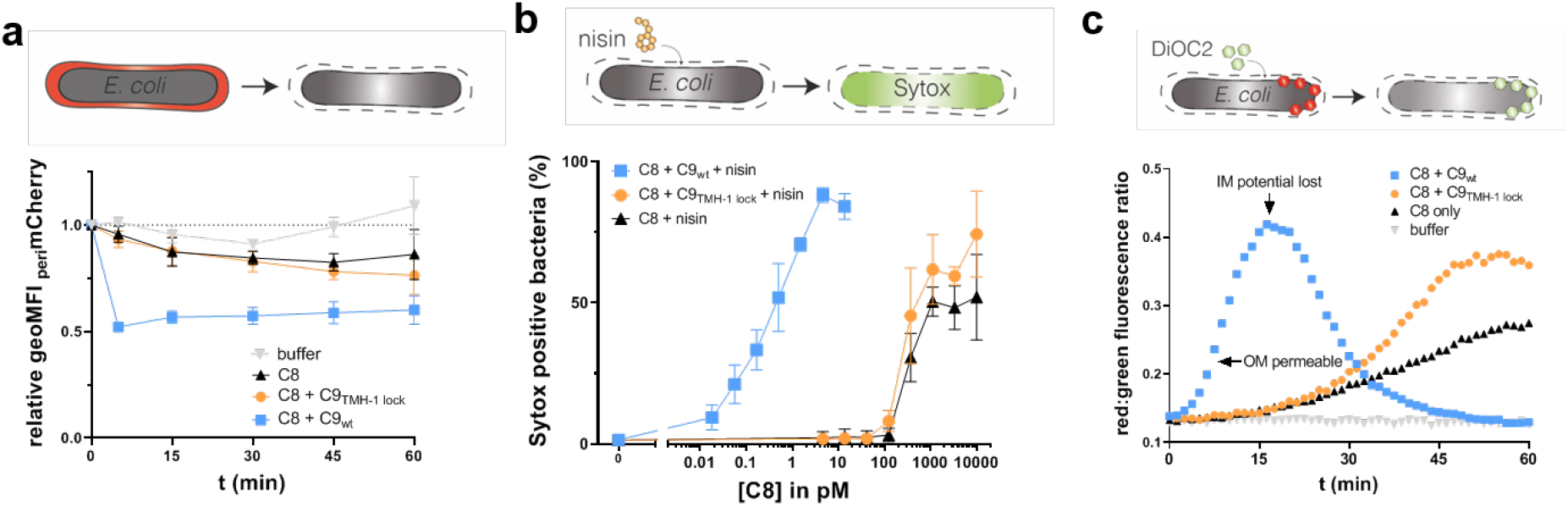
Polymerization of C9 increases OM damage by MAC pores. *E. coli* MG1655 were labelled with C5b-7 by incubating them in 10% C8-depleted serum for 30 minutes. Bacteria were washed and next incubated with buffer, 10 nM C8 with 20 nM C9_wt_ or C9_TMH-1 lock_ to measure damage to the bacterial OM. a) periplasmic mCherry (_peri_mCherry, red) leakage was measured at different time points by flow cytometry and represented as relative _peri_mCherry fluorescence compared to t=0. b) C5b-7 labelled bacteria were incubated with a concentration range of C8 and 20 nM C9_wt_ or C9_TMH-1 lock_ supplemented with 3 μg/ml nisin for 30 minutes. Nisin influx through OM was measured using Sytox to determine the percentage of cells that have a damaged bacterial IM as read-out for bacterial killing by flow cytometry after 30 minutes. c) DiOC_2_ enters the cells when the OM is permeable and shifts from green to red fluorescence in cells with an intact inner membrane (IM) potential. DiOC_2_ shifts back to green fluorescence when the IM potential is lost. Flow cytometry data are represented by geoMFI values of the bacterial population. Data represent mean +/− SD of three independent experiments. Multiwell plate-reader assays are shown by one representative experiment that has been repeated at least three times.

We also assessed if C9 polymerization affects the influx of extracellular molecules through the OM. We have previously shown that perturbation of the OM by MAC pores can sensitize Gram-negative bacteria to the antibiotic nisin (3.4 kDa), which normally cannot pass through the OM of Gram-negative bacteria [26]. A 1,000-fold more C5b-8 was required to sensitize bacteria to nisin with C9_TMH-1 lock_ compared to C9_wt_ (**Fig. 3b**), suggesting that polymerization of C9 strongly enhanced damage to the OM. Finally, we measured the influx of DiOC_2_ (0.5 kDa) through the OM in the presence or absence of polymerization. DiOC_2_ is a green fluorescent dye that shifts to red fluorescence when it is incorporated in membranes with a membrane potential, which in the case of *E. coli* is the IM. With C9_TMH-1 lock_, the increase in red:green fluorescence ratio was delayed compared to C9_wt_, suggesting that polymerization of C9 enhanced the influx of DiOC_2_ through the OM (**Fig. 3c**). When bacteria die the IM potential is lost, which results in a drop of red fluorescence. Loss of IM potential, initiated at the peak in red:green ratio, was also delayed with C9_TMH-1 lock_ compared to C9_wt_ (**Fig. 3c**). Interestingly, C9_TMH-1 lock_ did cause more rapid influx of DiOC_2_ (**Fig. 3c**) and nisin (**S2**) through the OM compared to C5b-8 alone. This also suggests that binding of C9 to C5b-8 in the absence of polymerization slightly increases OM damage. Altogether, these data highlight that polymerization of C9 increases damage to the OM.

### Polymerization of C9 is rate-limiting in the assembly of complete MAC pores

A recent study suggested that binding of the first C9 to C5b-8 is a kinetic bottleneck in the formation of a complete MAC pore on model lipid membranes [30]. Here, we used C9_TMH-1 lock_ to study if this is also the case when MAC pores assemble on bacteria. Therefore, *E. coli* MG1655 were labelled with a constant amount of C5b-8 while the amount of available C9_wt_ and C9_TMH-1 lock_ was limited. Binding of C9_wt_ was comparable to C9_TMH-1 lock_ when the amount of available C9 was below 1 nM (**Fig. 4a**), suggesting that there is no polymerization when the amount of available C9 is limited. As the amount of available C9 was increased, binding of C9_wt_ continued to increase up to 10-fold higher than C9_TMH-1 lock_ (**Fig. 4a**), whereas binding of C9_TMH-1 lock_ saturated (**Fig. 4a**). This suggests that C9 starts polymerizing when all C5b-8 complexes have bound one C9 molecule. This was confirmed by antibody staining (**Fig. 4c**) and SDS-PAGE (**Fig. 4d**), showing mainly mono-C9 when C9 is limited and the appearance of poly-C9 at C9 concentrations above 1 nM. Moreover, when the amount of surface-bound C5b-8 was varied by titrating C8, the difference in C9_wt_ binding compared to C9_TMH-1 lock_ binding decreased when an excess of C8 was added (**S3a,b**). _peri_mCherry leakage (**S3c**) and SDS-PAGE (**S3d**) confirmed that an excess of C8 decreased the relative abundance of poly-C9 compared to mono-C9. Altogether, these data suggest that polymerization of C9 is rate-limiting in the assembly of complete MAC pores on bacteria.

**Figure 4.**
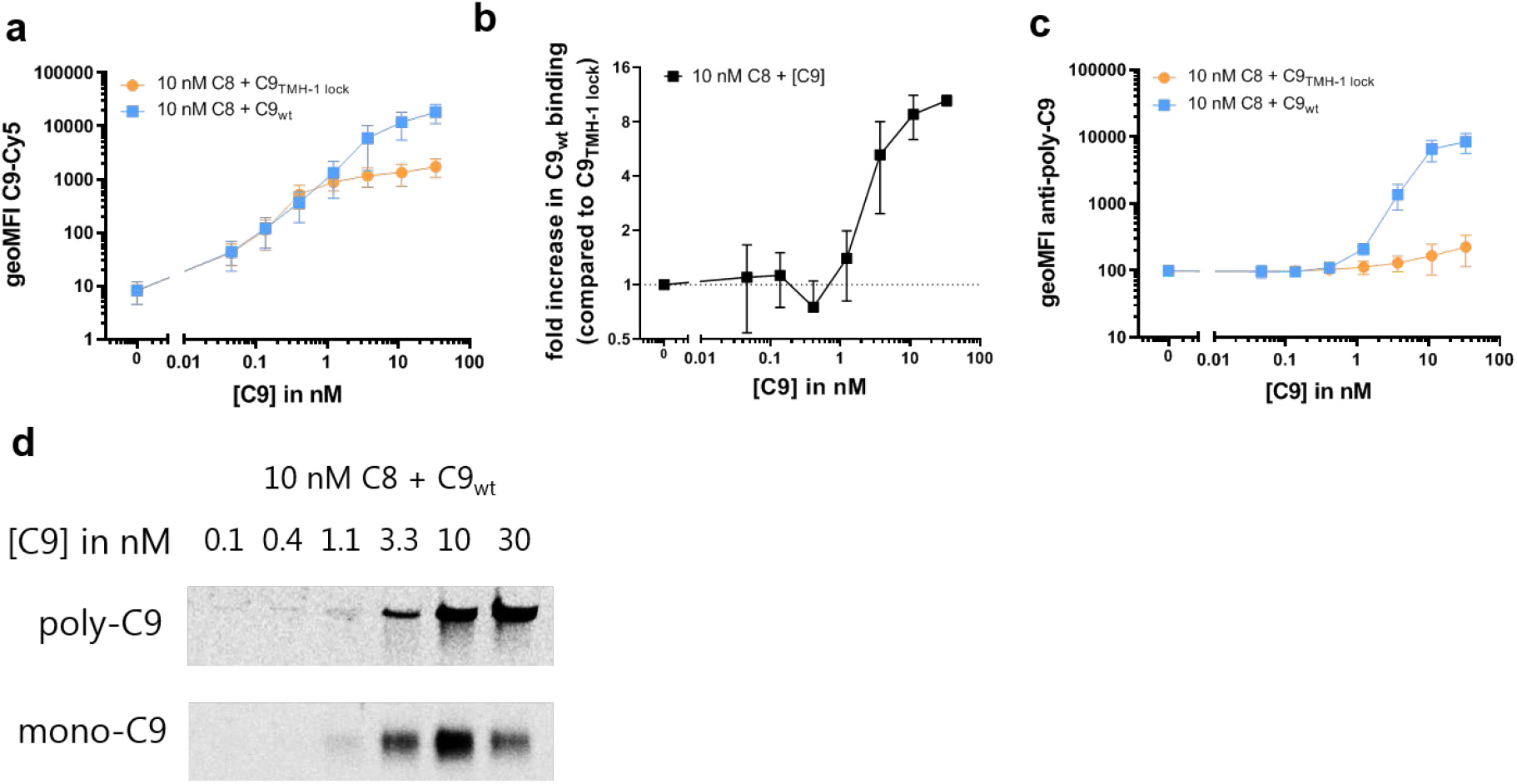
Polymerization of C9 is rate-limiting in the assembly of a complete MAC pore. *E. coli* MG1655 were labelled with C5b-7 by incubating them in 10% C8-depleted serum for 30 minutes. Bacteria were washed and next incubated with 10 nM C8 and a concentration range of Cy5-labelled C9_wt_ or C9_TMH-1 lock_ for 30 minutes to measure binding of C9. a) Binding of Cy5-labelled C9_wt_ or C9_TMH-1 lock_ to bacteria measured by flow cytometry. b) The relative increase in C9_wt_ binding compared to bacteria labelled with C9_TMH-1 lock_ was calculated as indication for C9 polymerization. c) Bacteria were stained with AF488-labelled mouse anti-poly-C9 aE11-antibody and staining was measured by flow cytometry. d) Bacterial cell pellets were analyzed by SDS-PAGE for in-gel fluorescence of Cy5-labelled C9_wt_ to distinguish monomeric-C9 (mono-C9) from polymeric-C9 (poly-C9). Flow cytometry data are represented by geoMFI values of the bacterial population. Data represent mean +/− SD of three independent experiments. SDS-PAGE images are representative for at least three independent experiments.

### Polymerization of C9 enhances bacterial cell envelope damage and killing in serum

So far, we have looked at the effect of C9 polymerization on the MAC in the absence of other serum components. Next, we wanted to see if polymerization of C9 also enhances bacterial cell envelope damage and killing in a serum environment. MG1655 was incubated in 3% C9-depleted serum with C9_wt_ or C9_TMH-1 lock_ and influx of DiOC_2_ or Sytox were measured over time in a plate-reader to measure OM and IM damage respectively. OM damage (**Fig. 5a**) preceded IM damage (**Fig. 5b**), but both were delayed by 30 minutes with C9_TMH-1 lock_ compared to C9_wt_. Also, bacterial viability was a 1,000-fold higher for C9_TMH-1 lock_ compared to C9_wt_ after 30 minutes (**Fig. 5c**). These data suggest that polymerization of C9 enhances bacterial cell envelope damage and killing in serum.

**Figure 5.**
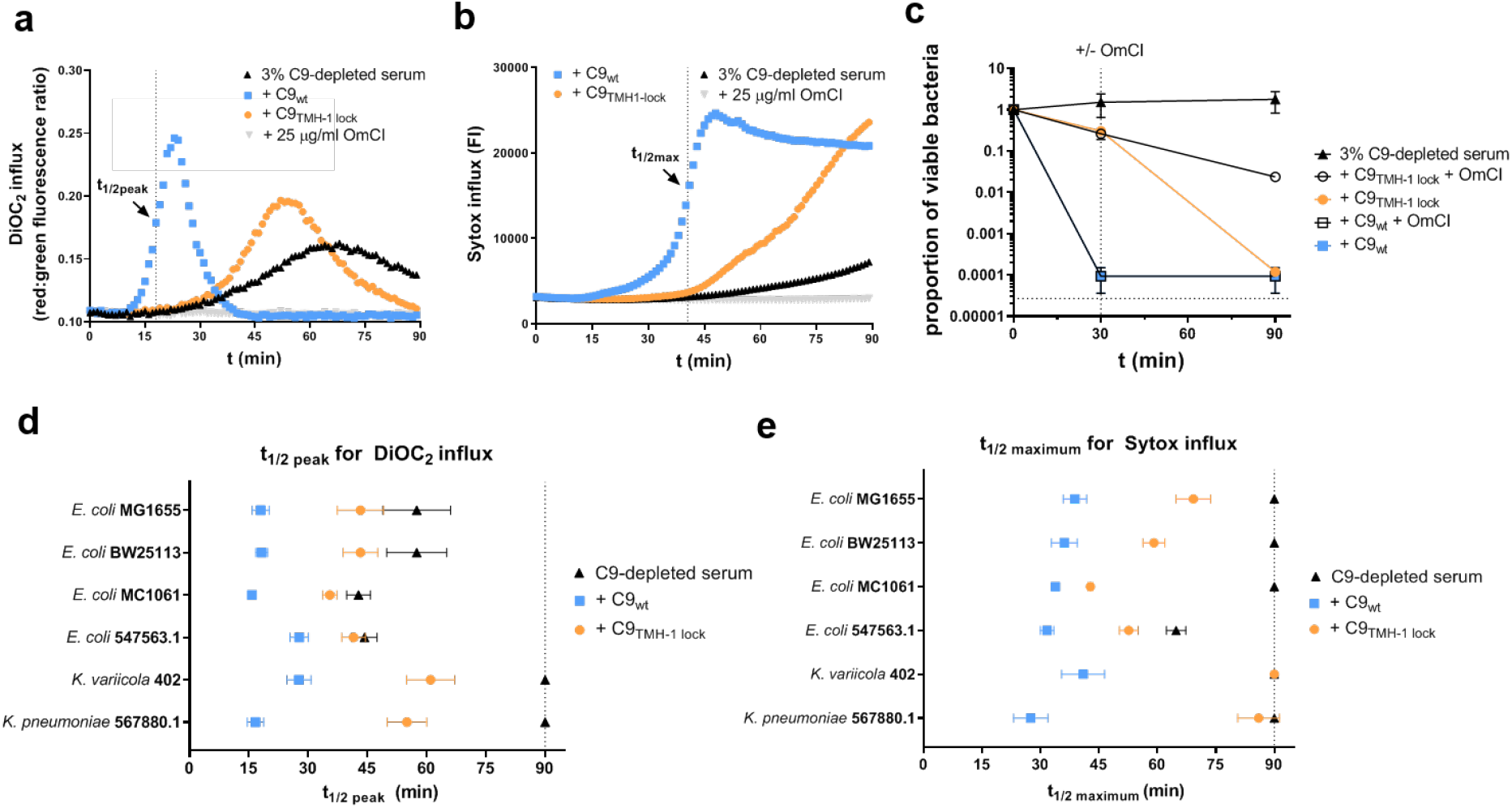
Polymerization of C9 enhances bacterial cell envelope damage and killing in serum. *E. coli* MG1655 were incubated in 3% C9-depleted serum supplemented with a physiological concentration (= 25 nM) of C9_wt_ or C9_TMH-1 lock_ for 90 minutes. As negative control, 25 μg/ml C5 conversion inhibitor OmCI was added to the serum. a) OM damage was measured by DiOC_2_ influx, which was determined by the shift in red:green fluorescence ratio over time in a multiwell plate-reader assay. b) IM damage was measured by Sytox influx over time in a multiwell plate-reader assay. c) Bacterial viability was determined at different time points by counting colony forming units (CFU’s) and calculating the proportion of viable cells compared to t=0. At t=30 (vertical dotted line), buffer was added (closed symbols) or OmCI (open symbols) to stop MAC formation. The horizontal dotted line represents the detection limit of the assay. d-e) Other *E. coli* strains (BW25113, MC1061, 547563.1) and *K. variicola* 402 were also incubated in 3% C9-depleted serum supplemented with C9_wt_ or C9_TMH-1 lock_ for 90 minutes. *K. pneumoniae* 567880.1 was incubated in 10% C9-depleted serum because of less efficient activation of the complement cascade and was therefore also supplemented with 80 nM C9_wt_ or C9_TMH-1 lock_. d) OM damage was represented for all strains by the time when DiOC_2_ influx reached half the value of the peak (t_1/2peak_, shown in a for C9_wt_). e) IM damage was represented for all strains by the time when Sytox fluorescence reached half the maximum value (t_1/2maximum_, shown in b for C9_wt_). Multiwell plate-reader assays (a, b) are shown by one representative experiment that has been repeated at least three times. Data represent mean +/− SD of three independent experiments (c, d and e).

Interestingly, bacteria were killed with C9_TMH-1 lock_ to a comparable level as C9_wt_ after 90 minutes. This was not observed with C9_TMH-1 lock_ in the absence of serum (**S4a**), which indicated that the presence of serum affects killing by the MAC, even in the absence of polymerized C9. In serum, complement activation is still ongoing, which could ultimately result in more MAC on the surface of bacteria and explain why viability still decreases in time. Indeed, stopping further MAC formation, by adding C5 conversion inhibitor OmCI after 30 minutes, prevented bacterial killing in C9-depleted serum with C9_TMH-1 lock_ over a 100-fold (**Fig. 5c**). Serum also contains bactericidal enzymes that can more efficiently pass through a damaged OM, such as lysozyme (14.7 kDa) and type IIa secreted phospholipase 2A (PLA, 14.5 kDa) [31]. Both lysozyme and PLA decreased bacterial viability when C5b-8 labelled bacteria were incubated with C9_TMH-1 lock_ compared to C5b-8 alone (**S4b**), suggesting that these serum proteins also contribute to bacterial killing in serum in the absence of polymerized C9. Nonetheless, taken together our data mainly highlight that polymerization of C9 enhances the efficiency by which MAC pores damage the bacterial cell envelope and kill bacteria in serum.

### Polymerization of C9 enhances cell envelope damage for several *E. coli* and *Klebsiella* strains in serum

We wondered if our findings on *E. coli* MG1655 could also be extrapolated to other *E. coli* strains and other Gram-negative species. First, the effect of C9 polymerization on OM and IM damage was measured for three other complement-sensitive *E. coli* strains (BW25113, MC1061 and 547563.1) in serum. To compare strains, time points were interpolated at which the red:green fluorescence ratio of DiOC_2_ reached half the value of the peak (t_1/2peak_, shown in **Fig. 5a**) as measure for OM damage, and at which half the maximum Sytox value was reached (t_1/2maximum_, shown in **Fig. 5b**) as measure for IM damage. Both OM damage (**Fig. 5d, S5a**) and IM damage (**Fig. 5e, S5b**) were delayed with C9_TMH-1 lock_ compared to C9_wt_ for all tested *E. coli* strains. C9_TMH-1 lock_ did increase OM damage and IM damage compared to C9-depleted serum alone for three out of four *E. coli* strains (**Fig. 5d,e and S5a,b**). For one strain, 547563.1, C9_TMH-1 lock_ only slightly enhanced IM damage, since C9-depleted serum alone already damaged the IM of these bacteria (**Fig. 5e, S5b**). OM damage (**Fig. 5d, S5a**) and IM damage (**Fig 5e, S5b**) were also delayed with C9_TMH-1 lock_ compared to C9_wt_ for two clinical *Klebsiella* isolates (*Klebsiella variicola* 402 and *Klebsiella pneumoniae* 567880.1). C9_TMH-1 lock_ did not cause IM damage within 90 minutes for *Klebsiella,* suggesting that for *Klebsiella* IM damage by MAC pores was more dependent on polymerization of C9 than for *E. coli*. Altogether, these data suggest that C9 polymerization enhances cell envelope damage for multiple complement-sensitive *E. coli* and *Klebsiella* strains in serum.

### Polymerization of C9 is impaired on complement-resistant *E. coli* strains

In our previous study we have identified *E. coli* isolates that survive killing by MAC pores because MAC does not stably insert into the OM [15]. Here, we wanted to see if C9 polymerizes on these complement-resistant *E. coli* strains (clinical isolates 552059.1, 552060.1 and 567705.1). All three complement-resistant strains express LPS O-Ag, whereas only one out of four complement-sensitive strains used in this study did as well (**S6a**). Binding of C9_TMH-1 lock_ in 3% C9-depleted serum was comparable between all strains, suggesting that the total amount of C5b-8 on the surface was comparable for complement-sensitive and complement-resistant *E. coli* (**Fig. 6a**). However, for complement-resistant *E. coli* binding of C9_wt_ was similar to C9_TMH-1 lock_, suggesting that C9 did not polymerize (**Fig. 6b**). By contrast, binding of C9_wt_ was 5-to 15-fold higher than C9_TMH-1 lock_ for all four complement-sensitive *E. coli* (**Fig. 6b**). Adding up to 10-fold more C9 slightly increased the difference between C9_wt_ binding and C9_TMH-1 lock_ 2- to 3-fold on complement-resistant 552059.1 (**Fig. 6c**), but this difference was still 5-fold lower compared to complement-sensitive MG1655. Although this suggested that some C9 polymerized on 552059.1, bacteria were still not killed (**S6b**). These data suggest that polymerization of C9, but not binding of C9 to C5b-8, is impaired on complement-resistant *E. coli*.

**Figure 6.**
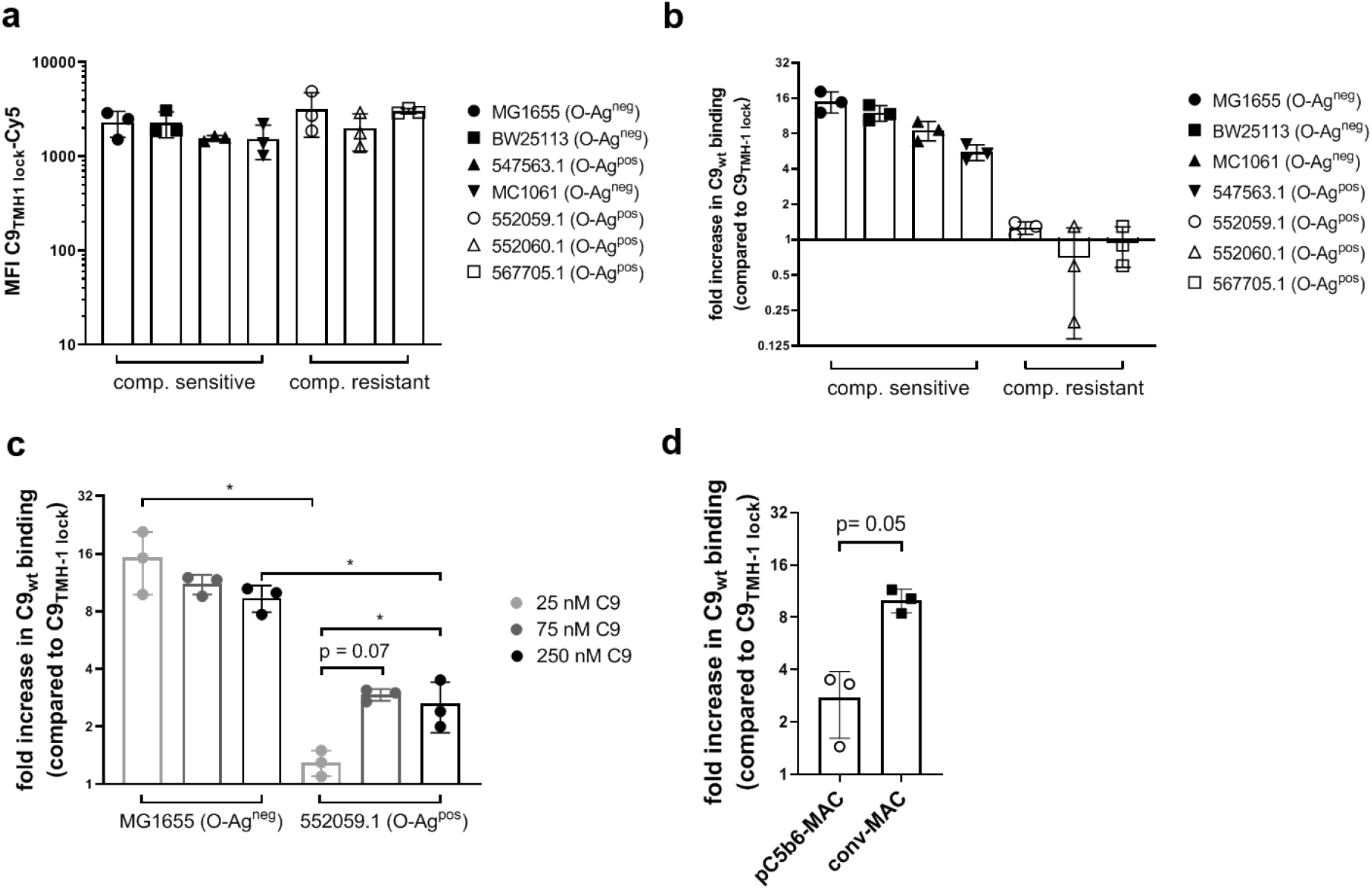
Polymerization of C9 is impaired on complement-resistant *E. coli* strains. Complement-resistant (552059.1, 552060.1 and 567705.1) and complement-sensitive (MG1655, BW25113, MC1061 and 547563.1) *E. coli* strains were incubated in 3% C9-depleted serum supplemented with a physiological concentration (= 25 nM) of Cy5-labelled C9_wt_ or C9_TMH-1 lock_ for 30 minutes to measure C9 binding by flow cytometry. a) Binding of Cy5-labelled C9_TMH-1 lock_ to bacteria. b) The relative increase in C9_wt_ binding compared to bacteria labelled with C9_TMH-1 lock_ was calculated as indication for C9 polymerization. c) The relative increase in C9_wt_ binding compared to bacteria labelled with C9_TMH-1 lock_ was compared for complement-sensitive MG1655 and complement-resistant 552059.1 at different C9 concentrations. d) MG1655 was labelled with convertases in 10% C5-depleted serum. Next, bacteria were washed and 3 nM of preassembled C5b6 (pC5b6) or C5 and C6 (conv-MAC) were added, together with excess 20 nM C7, 20 nM C8 and 50 nM Cy5-labelled C9_wt_ or C9_TMH-1 lock_ for 30 minutes. The relative increase in C9_wt_ binding compared to bacteria labelled with C9_TMH-1 lock_ was represented. Flow cytometry data are represented by MFI values of the bacterial population. Data represent individual values with mean +/− SD of three independent experiments. Statistical analysis was done on ^2^log-transformed data (a, c, d) using a paired one-way ANOVA with Tukey’s multiple comparisons’ test. Significance was shown as * p ≤ 0.05.

Finally, our previous study also indicated that although C5b-7 binds to these complement-resistant *E. coli*, it does not properly anchor to the OM [15]. This improper anchoring of C5b-7 ultimately resulted in instable insertion of the MAC, which could be mimicked on complement-sensitive *E. coli* using pC5b6 to assemble MAC [14,15]. In short, MG1655 were labelled with convertases in 10% C5-depleted serum, washed and afterwards pC5b6 was added in combination with C7, C8 and C9_wt_ or C9_TMH-1 lock_. For this pC5b6-MAC, C9_wt_ binding was 2- to 3-fold higher compared to C9_TMH-1 lock_ (**Fig. 6c**), resembling the difference in binding seen for complement-resistant *E. coli* 552059.1. By contrast, when convertases on MG1655 converted C5 and directly assembled MAC pores (conv-MAC), C9_wt_ binding was 12-fold higher compared to C9_TMH-1 lock_ (**Fig. 6c**). Altogether, these data suggest that improper anchoring of C5b-7 results in impaired polymerization of C9 on complement-resistant *E. coli*.

## Discussion

Understanding how MAC pores damage the bacterial cell envelope and kill bacteria is important to understand how the complement system prevents infections. Although it was still unclear whether a completely assembled MAC pore is needed to kill bacteria, we here show that it is important to efficiently damage the bacterial cell envelope and rapidly kill multiple Gram-negative bacterial strains and species.

Our findings suggest that bacteria are killed more rapidly when C9 polymerizes, both by MAC alone as well as in a serum environment. Previous reports have already suggested that the absence of C9 delays bacterial killing [32], and have correlated the presence of polymeric-C9 to bacterial killing [33,34]. Our study extends on these insights, since it provides direct evidence that polymerization of C9 enhances bacterial killing by using a system in which C9 can bind to C5b-8 without polymerizing. Although Spicer *et al.* had already suggested that locking the TMH-1 domain of C9 could prevent polymerization of C9, we here validate that this does not affect binding of C9 to C5b-8 [27]. Apart from bacterial killing, our study also suggests that polymerization of C9 more efficiently damages both the OM and IM of the bacterial cell envelope. Polymerization of C9 greatly enhanced passage of small molecules through the OM and was required for passage of periplasmic proteins through the OM. OM damage preceded IM damage, corresponding with our earlier findings [14]. Here, we add to these insights showing that OM damage also precedes the loss of IM potential, which is a widely accepted characteristic of cell death.

Based on our results, we therefore hypothesize that extensive OM damage by MAC pores is driving bacterial killing. Binding of C8 to C5b-7 allows passage of small molecules through the OM (**Fig 7-I**), but these lesions have minimal effect on the IM and bacterial viability. Binding of C9 to C5b-8 without polymerization of C9 slightly increases damage to both the OM and IM (**Fig 7-II**). Although some residual polymerization of C9_TMH-1 lock_ can never be fully excluded, our data do not show any direct evidence that C9_TMH-1 lock_ polymerized. Based on the structure of monomeric-C9 [27], it is highly unlikely that the other TMH domain (TMH-2) of C9 can insert into the membrane when the TMH-1 domain is locked. Binding of C9 to C5b-8 without polymerization of C9 could affect the stability by which C8 is inserted into the membrane, which ultimately could affect OM damage. Nonetheless, our data highlight that polymerization of C9 drastically enhanced the damage of both the OM and IM damage and rapidly killed bacteria (**Fig 7-III**). Our study therefore primarily emphasizes that polymerization of C9 strongly enhances bacterial cell envelope damage and killing.

**Figure 7.**
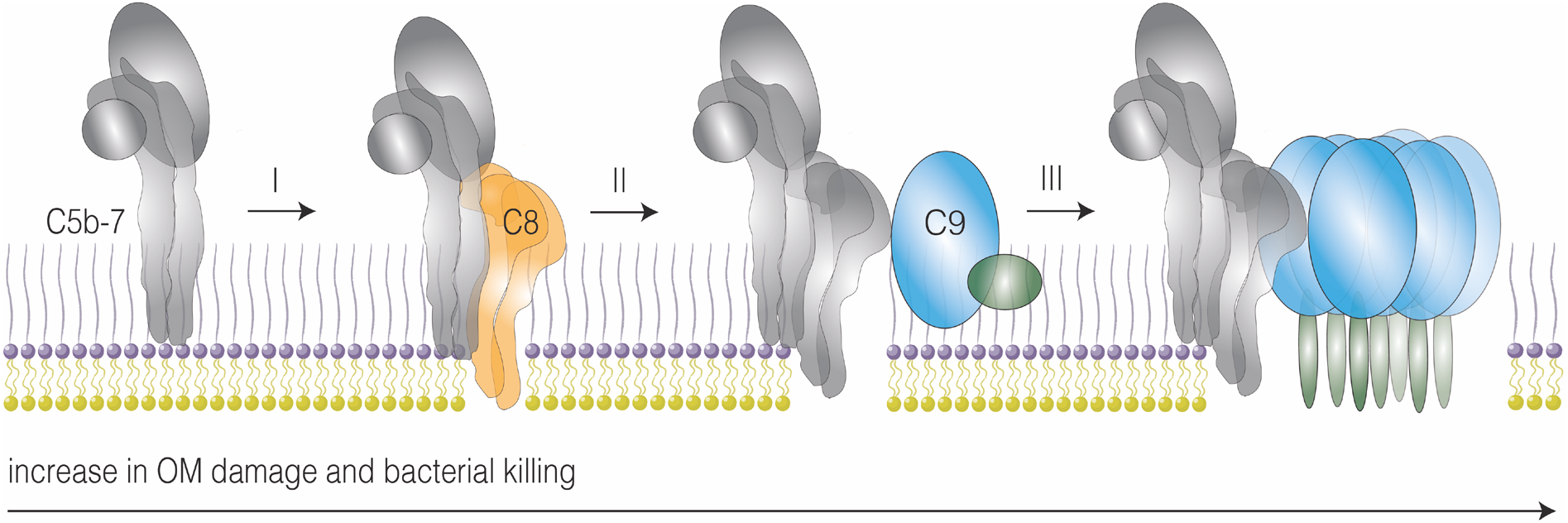
Assembly of complete MAC pores enhances OM damage and bacterial killing. Schematic overview of the assembly of complete MAC pores. I) C8 (orange) binds to membrane-anchored C5b-7 (grey) and subsequently inserts transmembrane β-hairpins into the bacterial outer membrane (OM), which causes small lesions in the OM. II) Binding of C9 (blue) to C5b-8 without polymerization of C9 slightly increases the OM damage and bacterial killing. III) Polymerization of C9 forms a transmembrane polymeric-C9 ring, which drastically increases OM damage and rapidly kills bacteria.

How OM damage by MAC pores destabilizes the IM and kills bacteria remains unclear. The OM is an essential load-bearing membrane that confers stability to the bacterial cell [35]. The extent of OM damage by MAC pores could therefore determine whether a cell can cope with the increase in osmotic pressure. OM damage by MAC pores could even directly interfere with osmoregulation of the cell, as has been suggested for antimicrobial peptides that damage the OM of *E. coli* [36]. Moreover, since metabolism and growth-phase have been associated with sensitivity to killing by MAC pores [37,38], OM damage could also induce a stress response that affects the capacity of a cell to survive envelope stress. C9-depleted serum alone damaged the IM for one *E. coli* strain, even though the amount of C5b-8, indicated by the binding of C9_TMH-1 lock_, and OM damage was comparable to other *E. coli* strains. On the other hand, for *Klebsiella* strains IM damage appeared more dependent on polymerization of C9 than for *E. coli* strains. These differences between strains and species could suggest that the capacity of a bacterium to cope with OM damage-related stress is a crucial determinant for bacterial killing by MAC pores. Further research looking directly at the cellular response of bacteria would be necessary to better understand how OM damage by MAC pores causes cell death.

Our data also suggest that polymerization of C9 is rate-limiting in the assembly of complete MAC pores on bacteria. Joiner *et al*. already demonstrated that limiting the C9 concentration decreased the ratio of C9:C7 on the surface [33]. We further extend on these insights, as our data also suggest that binding of C9 to C5b-8 is favoured over binding to an already unfurled C9 molecule in the nascent polymeric-C9 ring. By contrast, Parsons *et al*. [30] recently suggested that binding of C9 to C5b-8 is a kinetic bottleneck in the assembly of complete MAC pores. However, this study looked at MAC assembly in the presence of an excess of C9, which we here show to affect the assembly kinetics of MAC pores. Moreover, experiments in this study were done on model lipid-membranes that were not labelled with convertases, which also could have influenced the assembly of MAC pores [14].

Finally, our study highlights that polymerization of C9 is specifically impaired on *E. coli* that resist MAC-dependent killing. We have previously shown that C5b-7 could bind to these complement-resistant strains, but was improperly anchored to the OM which resulted in unstably inserted MAC [15]. Here, we extend on these findings showing that this improper anchoring of C5b-7 results in impaired polymerization of C9. How these complement-resistant *E. coli* affect the anchoring of C5b-7 and subsequent polymerization of C9 remains unclear. Polymerization of C9 correlated with the absence of LPS O-Ag in the *E. coli* strains used in this study. The presence and length of LPS O-Ag have been correlated with survival in serum before [23], and are thought to affect the distance of the nascent MAC to the OM and the accessibility of hydrophobic patches in the OM [8]. We therefore hypothesize that improper anchoring of C5b-7 initiates assembly of MAC further away from hydrophobic patches in the OM, which could prevent stable insertion of the nascent polymeric-C9 ring and result in an incomplete polymeric-C9 ring. Although LPS O-Ag modifications can also affect earlier steps in the complement cascade [9,10,39–41], this did not seem to play a role as the amount of C5b-8 on the surface was comparable between complement-sensitive and -resistant *E. coli*. Nonetheless, a more direct study of the effect of O-Ag on polymerization of C9 would be required to verify this hypothesis.

In conclusion, our study provides insight into how MAC pores damage the bacterial cell envelope and kill Gram-negative bacteria. Moreover, our study highlights how bacteria resist killing by MAC pores. These fundamental insights are important to understand how the complement system prevents infections and how bacteria escape killing by the immune system. Ultimately, these insights may guide the development of future immune therapy against bacteria.

## Materials & Methods

### Serum and complement proteins

Serum depleted of complement components C5, C8 or C9 was obtained from Complement Technology. Serum was thawed and aliquoted, but not subjected to any further freeze-thaw cycles. Preassembled C5b6 (pC5b6) and C8 were also obtained from Complement Technology. His-tagged complement components C5, C6 and C7 were expressed in HEK293E cells at U-Protein Express as described previously [15]. OmCI was produced in HEK293E cells at U-Protein Express as well and purified as described before [42]. Monoclonal mouse-anti poly-C9 (maE11, kindly provided by T. Mollness and P. Garred) was randomly labelled with NHS-Alexa Fluor AF488 (Thermofisher) according to manufacturer’s protocol. Lysozyme was obtained from Raybio and recombinant type IIa secreted phospholipase 2A (PLA) was kindly provided by Gérard Lambeau [43].

### Expression and purification of C9

C9_wt_ and C9_TMH-1 lock_ (F262C V405C) were cloned into the vector pcDNA34 (Thermo Fisher Scientific) that was modified with an NheI/NotI multiple cloning site. This vector contains a cystatin-S signal peptide and was further modified to encode for the expression of a C-terminal AAA-3x(GGGGS)-LPETGG-HHHHHH tag. gBlocks (Integrated DNA Technologies) containing codon optimized C9 sequences were cloned via Gibson assembly into the NheI/NotI digested pcDNA34 and transformed into Top10F *E. coli*. After verification of the correct sequence, the plasmids were used to transfect EXPI293F cells. EXPI293F cells were grown in EXPI293 medium (Life Technologies) in culture filter cap Erlenmeyer bottles (Corning) on a rotation platform (125 rotations/min) at 37°C, 8% CO_2_. One day before transfection, cells were diluted to 2×10^6^ cells/ml. The next day, cells were diluted to 2×10^6^ cells/ml using SFM4Transfx-293 medium, containing UltraGlutamine I (VWR International) prior to transfection using PEI (Polyethylenimine HCl MAX; Polysciences). 0.5 μg DNA/ml cells, containing 50% empty (dummy) vector, was added to Opti-MEM (1:10 of total volume; Gibco) and gently mixed. After adding 1 μg/ml PEI in a PEI/DNA (w/w) ratio of 5:1, the mixture was incubated at room temperature for 20 min and added dropwise to cells while manually rotating the culture flask. After 3.5 hours, 1 mM valproic acid (Sigma) was added. After 5 days of expression, the cell supernatant was collected by centrifugation and filtration (0.45 μm). Cell supernatant was diafiltrated over a 30 kDa membrane on a Quixstand (GE healthcare) to Tris/NaCl buffer (50 mM Tris/500 mM NaCl at pH 8.0). Proteins were finally loaded on a HisTrap HP Chelating column (GE healthcare) in Tris/NaCl buffer supplemented with 40 mM imidazole and eluted with 150 mM imidazole. Final purification was done by size-exclusion chromatography (SEC) on a Superdex 200 Increase column (GE Healthcare) on an Akta Explorer (GE Healthcare) with PBS. The concentration of proteins was determined by measuring absorbance at 280 nm and verified by SDS-PAGE.

### Site-specific fluorescent labelling of MAC components

C6 and C9 were labelled with fluorescent probes as described previously [14,15]. 50 μM of protein with C-terminal LPETGG-His tag was incubated with 25 μM His-tagged sortase-A7+ [44] and 1 mM GGG-substrate in Tris/NaCl buffer (50 mM Tris/300 mM NaCl at pH 7.8) for two hours at 4 °C. GGGK-FITC (Isogen Life Science) was used for C6-LPETGG-His and GGGK-azide (Genscript) for C9-LPETGG-His. Sortagged proteins were purified on a HisTrap FF column (GE Healthcare), which captures protein that was not sortagged and still contains a His-tag. FITC-labelled C6 was directly purified by SEC on a Superdex 200 Increase column on the Akta Explorer with PBS. GGG-azide labelled proteins were concentrated to 25 μM on a 30 kDa Amicon Tube (Merck Millipore) in Tris/NaCl buffer and next labelled with 100 μM DBCO-Cy5 (Sigma Aldrich) via copper-free click chemistry for 3h at 4 °C. Finally, Cy5-labelled proteins were also purified by SEC on a Superdex 200 Increase column with PBS. Labelling of the proteins was monitored during SEC by measuring absorbance at 280 nm (protein), 488 nm (FITC) and 633 nm (Cy5) nm and finally verified by SDS-PAGE by measuring in-gel fluorescence with LAS4000 Imagequant (GE Healthcare).

### Bacterial strains

Unless otherwise specified, the common laboratory *E. coli* strain MG1655 was used in our experiments. For experiments where leakage of periplasmic mCherry was measured, MG1655 was used transformed with pPerimCh containing a constitutively expressed periplasmic mCherry (_peri_mCherry) previously used in [14]. Other laboratory *E. coli* strains that were used in this study included BW25113 and MC1061. Clinical isolates, namely *E. coli* 547563.1, 552059.1, 552060.1, 567705.1, *Klebsiella variicola* 402 and *Klebsiella pneumonia*e 567880.1, were obtained from the clinical Medical Microbiology department at the University Medical Center Utrecht.

### Bacterial growth

For all experiments, bacteria were plated on Lysogeny Broth (LB) agar plates. Single colonies were picked and grown overnight at 37 °C in LB medium. For MG1655 transformed with pPerimCh, LB was supplemented with 100 μg/ml ampicillin. The next day, subcultures were grown by diluting at least 1/30 and these were grown to mid-log phase (OD600 between 0.4 – 0.6). Once grown to mid-log phase, bacteria were washed by centrifugation three times (11000 rcf for 2 minutes) and resuspended to OD 1.0 (~1 × 10^9^ bacteria/ml) in RPMI (Gibco) + 0.05% human serum albumin (HSA, Sanquin).

### Complement labelling and serum bactericidal assays

For MAC-specific bactericidal assays, bacteria were labelled with C5b-7 as described previously [15]. In short, bacteria (~1 × 10^8^ bacteria/ml) were incubated with 10 % C8-depleted serum for 30 minutes at 37 °C, washed three times and resuspended in RPMI-HSA. C5b-7 labelled bacteria (~5 × 10^7^ bacteria/ml) were incubated for 30 minutes at 37 °C with 10 nM C8 and 20 nM C9, unless stated differently. When C9 was reduced with 10 mM dithiothreitol (DTT), 20 nM C8 was added to bacteria (~5 × 10^7^ bacteria/ml) 15 minutes at RT before C9 was added to allow binding of C8 to C5b-7. To label bacteria with convertases, bacteria (~1 × 10^8^ bacteria/ml) were incubated with 10 % C5-depleted serum for 30 minutes at 37 °C as described previously [14,15]. Washing steps were done by pelleting bacteria at 11,000 rcf for 2 minutes and washing with RPMI-HSA. For serum bactericidal assays, bacteria (~5 × 10^7^ bacteria/ml) were incubated with 3% C9-depleted serum supplemented with physiological concentrations of C9 (100% serum ± 1 μM) for 30 minutes at 37 °C, unless stated differently. KP880.1 was incubated in 10% C9-depleted serum with the corresponding physiological concentration of C9. Blocking of C5 conversion in serum was done with 25 μg/ml OmCI as final concentration. For assays where nisin (Handary, SA, Brussels) was added, 3 μg/ml was used as final concentration.

### Flow cytometry

Complement-labelled bacteria (~5 × 10^7^ bacteria/ml) were incubated with 2.5 μM of Sytox Blue Dead Cell stain (Thermofisher). Samples were diluted to ~2.5 × 10^6^ bacteria/ml in RPMI-HSA and subsequently analyzed on a MACSquant VYB (Miltenyi Biotech) for Sytox and _peri_mCherry fluorescence. Poly-C9 deposition was measured by incubating bacteria (2.5 × 10^7^ bacteria/ml) with 6 μg/ml monoclonal AF488 labelled mouse-anti poly-C9 (maE11) for 30 minutes at 4 °C. For C9 binding in serum, bacteria were stained with 1 μM Syto9 (Thermofisher) to exclude serum noise events. Samples were next diluted to ~2.5 × 10^6^ bacteria/ml in 1.1% paraformaldehyde and subsequently analyzed on the BD FACSVerse flow cytometer for Cy5 and maE11-AF488 or Syto9 fluorescence. Flow cytometry data was analysed in FlowJo V.10. Bacteria were gated on forward scatter and side scatter. In serum, an additional trigger was placed on Syto9 fluorescence. Sytox positive cells were gated such that the buffer only control had <1 % positive cells.

### Poly-C9 detection by SDS-PAGE

Bacterial labelled with complement components were collected by spinning bacteria down 11,000 rcf for 2 minutes and subsequently washing cell pellets twice in RPMI-HSA. Cell pellets were resuspended and diluted 1:1 in SDS sample buffer (0.1M Tris (pH 6.8), 39% glycerol, 0.6% SDS and bromophenol blue) supplemented with 50 mg/ml DTT and placed at 95 °C for 5 minutes. Samples were run on a 4-12% Bis-Tris gradient gel (Invitrogen) for 75 minutes at 200V. Gels were imaged for 10 minutes with increments of 30 seconds on the LAS4000 Imagequant (GE Healthcare) for in-gel Cy5 fluorescence. Monomeric-C9 (mono-C9) and polymeric-C9 (poly-C9) were distinguished by size, since mono-C9 runs at 63 kDa and poly-C9 is retained in the comb of the gel.

### Bacterial viability assay

Bacteria were treated with MAC components or serum as described above. Next, colony forming units (CFU) were determined by making serial dilutions in PBS (100, 1.000, 10.000 and 100.000-fold). Serial dilutions were plated in duplicate on LB agar plates and incubated overnight at 37 °C. The next day, colonies were counted and the corresponding concentration of CFU/ml was calculated.

### Multi-well fluorescence assays

Bacteria (~5 × 10^7^ bacteria/ml) added to RPMI-HSA supplemented with 1 μM of Sytox Green Dead Cell stain (Thermofisher) or 30 μM DiOC_2_ (PromoCell). Bacteria were next incubated with MAC components, serum and/or nisin as described above. Fluorescence was measured every 60 seconds on a Clariostar platereader (BMG labtech). Sytox green fluorescence was measured using an excitation wavelength of 484-15 nm and emission wavelength of 527-20. DiOC_2_ fluorescence was measured using an excitation wavelength 484-15 nm and emission wavelength of 527-20 (green) and 650-24 (red). For DiOC_2_, the red fluorescence was divided by the green fluorescence to determine the red:green fluorescence ratio. t_1/2peak_ for DiOC_2_ was interpolated at half the value of the peak, which was calculated by substracting the background ratio value at t=0 from the peak ratio value and dividing by two. t_1/2maximum_ for Sytox was interpolated at half the value of the maximum Sytox value, which was calculated by substracting the fluorescence value at t=0 from the maximum fluorescence value and dividing by two.

### LPS O-antigen Silver staining

*E. coli* strains were typed for LPS O-Ag by Silver staining after SDS-PAGE based on [45,46]. In short, bacteria were scraped from blood agars plates in PBS and incubated at 56 °C for 60 minutes. Cell pellets were next deproteinated with 400 μg/ml proteinase K for 90 minutes and diluted in 2x Laemli buffer with 0.7 M beta-mercaptoethanol. Cell pellets were run on a 4-12% BisTris gel as described above and fixed overnight in fixing buffer (40% ethanol + 4% glacial acetic acid). The gel was oxidized for 5 minutes in fixing buffer supplemented with 0.6% periodic acid. The gel was then stained for 15 minutes with freshly prepared 0.3% silver nitrate in 0.125 M sodium hydroxide and 0.3% ammonium hydroxide. Finally, the gel was developed for 7 minutes in developer solution (0.25% citric acid + and 0.2% formaldehyde). In between steps, the gel was washed three times with MilliQ. Lipid A and LPS core were distinguished from LPS O-Ag based on size.

### Data analysis and statistical testing

Unless stated otherwise, graphs are comprised of at least three biological replicates. Statistical analyses were performed in GraphPad Prism 8 and are further specified in the figure legends.

## Acknowledgements

This work was funded by an ERC Starting grant (639209-ComBact, to S.H.M.R). The authors would like to acknowledge Lisanne de Vor and Myrthe Reiche for critically reading the manuscript.

## Author contributions

D.J.D., S.H.M. and B.W.B. conceived the project and designed the experiments. D.J.D. and C.J.C.H. performed protein purifications and fluorescent labelling. D.J.D. performed poly-C9 detection by SDS-PAGE. D.J.D., D.A.C.H., M.R. and B.W.B. performed flow cytometry, multiwell fluorescence and bacterial killing assays. P.C.A. performed LPS O-Ag typing with input from D.A.C.S. D.J.D., D.A.C.H., M.R., D.A.C.S., S.H.M., B.W.B. analyzed the data. D.J.D., S.H.M. and B.W.B. wrote the manuscript with input from D.A.C.H., D.A.C.S. All authors approved the final version of the manuscript.

