## Supplementary information for "Polymerization of C9 enhances bacterial cell envelope damage and killing by membrane attack complex pores"

**Supplementary figure 1 (S1)**

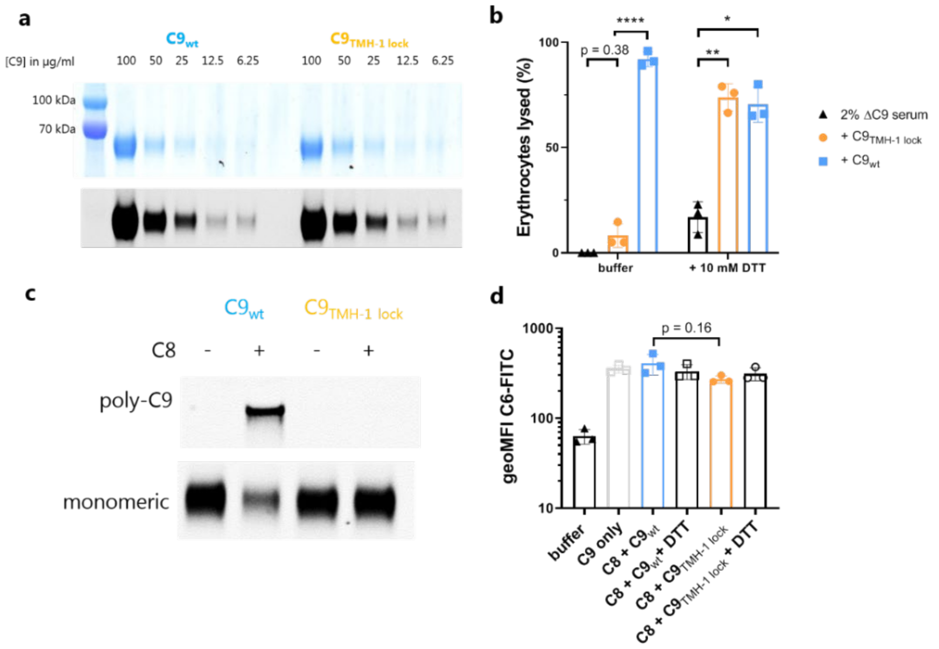


a) SDS-PAGE of a concentration range of Cy5-labelled C9$_{wt}$ and C9$_{TMH-1lock}$ (in µg/ml). Both InstantBlue staining (top) and in-gel Cy5 fluorescence (bottom) were shown. b) Sheep erythrocytes (4%) labelled with rabbit anti-sheep IgM were incubated in 2% C9-depleted serum for 30 minutes. Next, erythrocytes were washed and incubated with 30 nM C9$_{wt}$ or C9$_{TMH-1lock}$ in the presence or absence of 10 mM DTT. The percentage of lysed erythrocytes was calculated by adding MilliQ as 100% lysis or buffer as 0% lysis control after incubation with C9-depleted serum. c) 20 nM pC5b6, 20 nM C7, 20 nM C8 was incubated with 100 nM Cy5-labelled C9$_{wt}$ or C9$_{TMH-1lock}$. SDS-PAGE was done to distinguish monomeric-C9 (mono-C9) from polymeric-C9 (poly-C9) by in-gel Cy5 fluorescence. (d) C6-FITC binding to *E. coli* MG1655 measured by flow cytometry. Bacteria were labelled with C5b-7 by incubating them in 10% C8-depleted serum supplemented 12 µg/ml C6-FITC with for 30 minutes. Bacteria were washed and next incubated with 10 nM C8 for 15 minutes. Finally, 20 nM of Cy5-labelled C9$_{wt}$ or C9$_{TMH-1 lock}$ was added in the presence or absence of 10 mM DTT for 30 minutes. Flow cytometry data represent individual geoMFI values of the bacterial population. Graphs (b and d) represent three independent experiments with mean +/- SD. Statistical analysis was done using a paired one-way ANOVA with Tukey's multiple comparisons' test. For d, data were $^{10}$log-transformed. Significance was shown as * $p \leq 0.05$, ** $p \leq 0.005$, **** ** $p \leq 0.0001$

**Supplementary figure 2 (S2)**

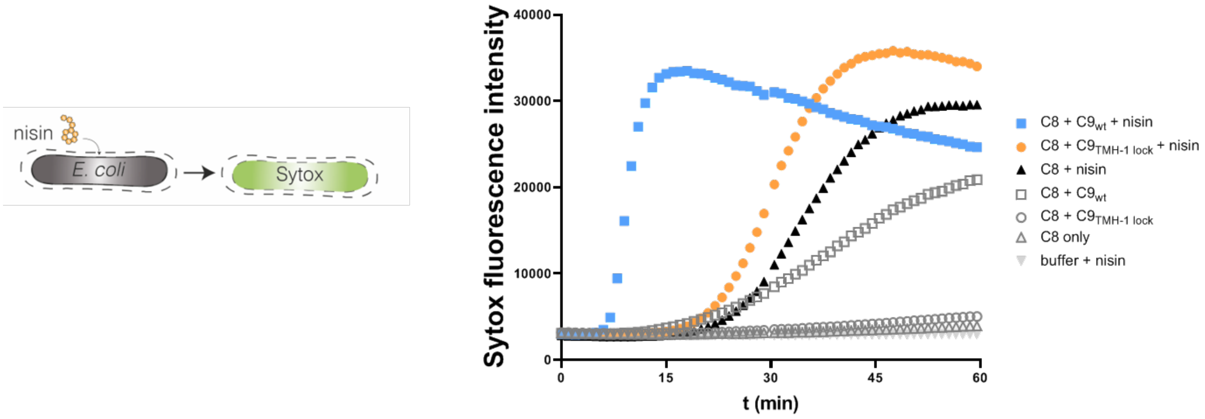


*E. coli* MG1655 were labelled with C5b-7 by incubating them in 10% C8-depleted serum for 30 minutes. Bacteria were washed and next incubated with 10 nM C8 and 20 nM of $C9_{wt}$ or $C9_{TMH\text{-}1\ lock}$ supplemented with or without 3 µg/ml nisin to measure passage of nisin through the OM. Nisin influx was determined by measuring Sytox influx over time in a multi-well plate-reader assay. One representative experiment was shown that has been repeated at least three times.

**Supplementary figure 3 (S3)**

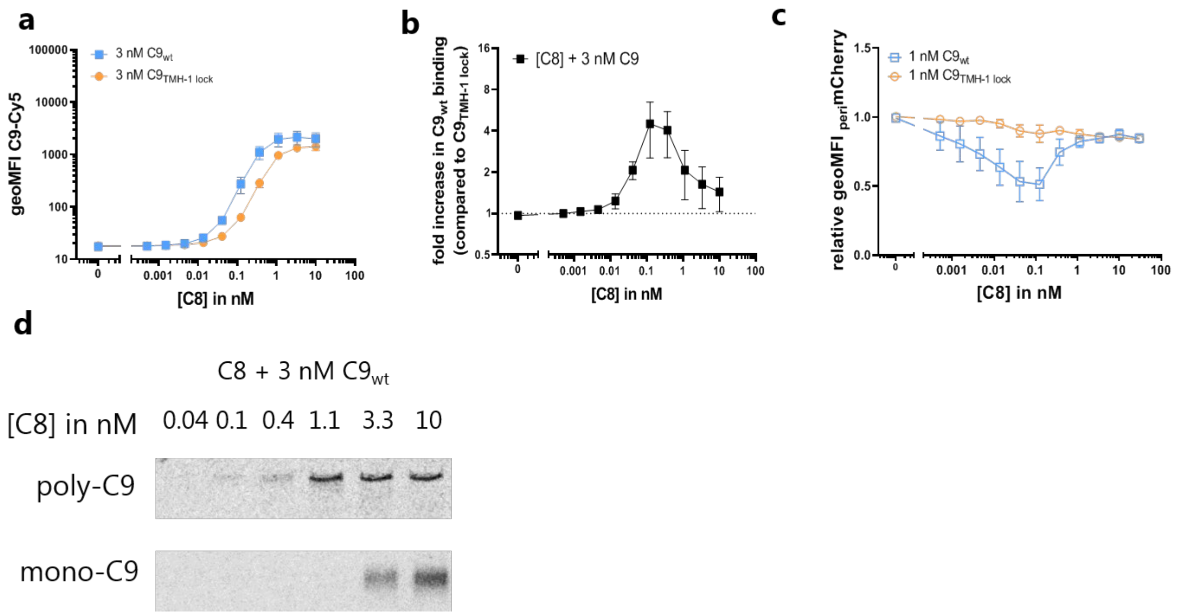


*E. coli* MG1655 were labelled with C5b-7 by incubating them in 10% C8-depleted serum for 30 minutes. Bacteria were washed and next incubated with a concentration of C8 and 3 nM (a,b,d) or 1 nM (c) Cy5-labelled $C9_{wt}$ or $C9_{TMH\text{-}1\ lock}$ for 30 minutes. a) Binding of Cy5-labelled $C9_{wt}$ or $C9_{TMH\text{-}1\ lock}$ to bacteria measured by flow cytometry. b) The relative increase in $C9_{wt}$ binding compared to bacteria labelled with $C9_{TMH\text{-}1\ lock}$ was calculated as indication for C9 polymerization. c) periplasmic mCherry ($_{peri}$mCherry) leakage was measured after 30 minutes by flow cytometry and represented as relative $_{peri}$mCherry fluorescence compared to t=0. d) Bacterial cell pellets were analyzed by SDS-PAGE for in-gel fluorescence of Cy5-labelled $C9_{wt}$ to distinguish monomeric-C9 (mono-C9) from polymeric-C9 (poly-C9). Flow cytometry data are represented by geoMFI values of the bacterial population. Data represent mean +/- SD of three independent experiments. SDS-PAGE images are representative for at least three independent experiments.

**Supplementary figure 4 (S4)**

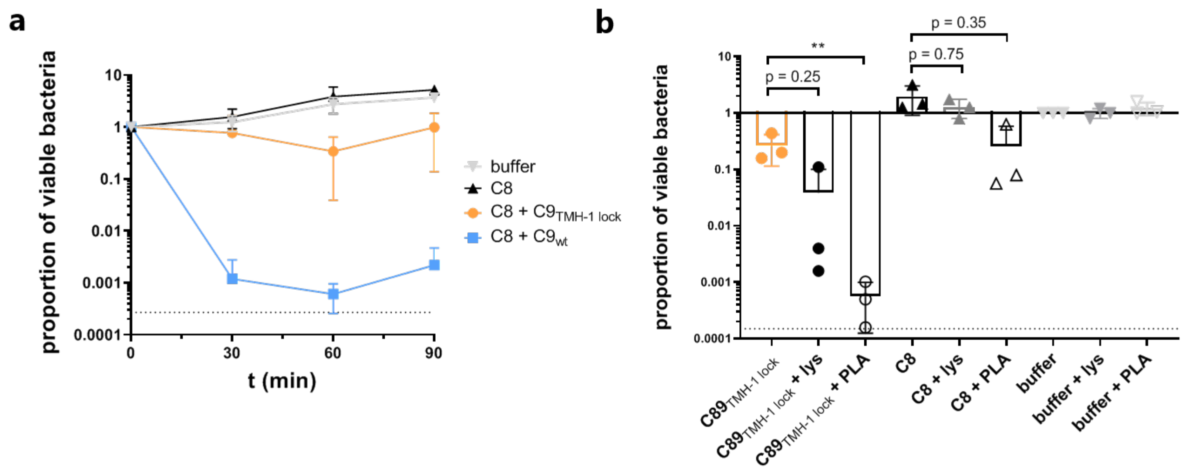


*E. coli* MG1655 were labelled with C5b-7 by incubating them in 10% C8-depleted serum for 30 minutes. Bacteria were washed and next incubated with 10 nM C8 and 20 nM of $C9_{wt}$ or $C9_{TMH\text{-}1\ lock}$. a) Bacterial viability was determined at different time points by counting colony forming units (CFU's) and calculating the proportion of viable cells compared to t=0. b) *E. coli* MG1655 bacteria were incubated with 10% C8 depleted serum for 30 minutes. After washing, bacteria were incubated with buffer, 10 nM C8 or 10 nM C8 + 20 nM C9 in the presence of 5 µg/ml lysozyme (lys) or 0.3 µg/ml recombinant type IIa secreted phospholipase 2A (PLA). Bacterial viability was determined after 90 minutes of incubation by counting CFU's and calculating the proportion of viable cells compared to t=0. b The horizontal dotted line represents the detection limit of the assay. Data represent mean +/- SD (a) or individual values with mean +/- SD of three independent experiments. Statistical analysis was done on $^{10}$log-transformed data (b) using a paired one-way ANOVA with Tukey's multiple comparisons' test. Significance was shown as ** $p \leq 0.005$.

### Supplementary figure 5 (S5)

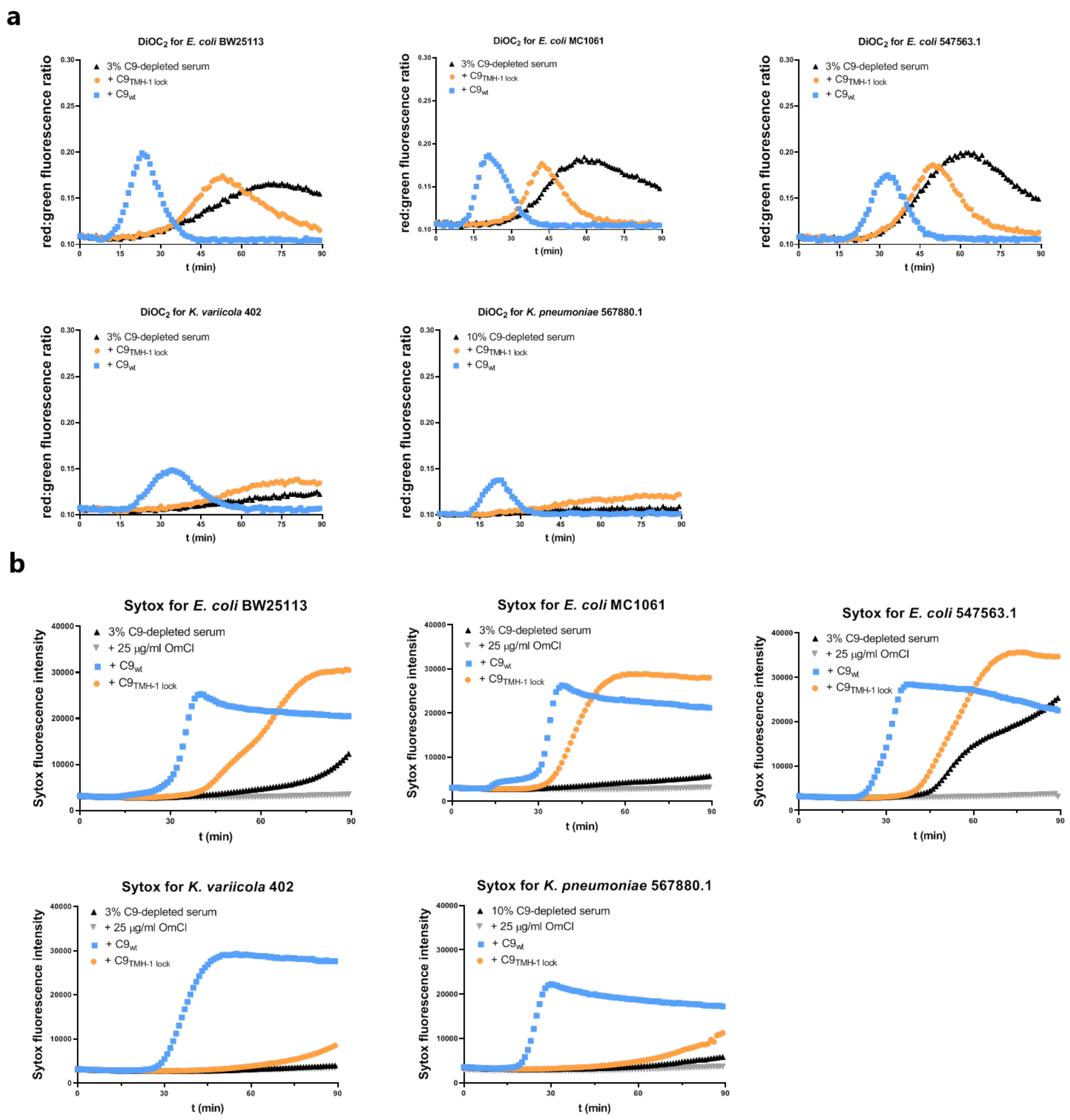


*E. coli* strains (BW25113, MC1061, 547563.1) and *K. variicola* 402 were incubated in 3% C9-depleted serum supplemented with a physiological concentration (= 25 nM) of $C9_{wt}$ or $C9_{TMH\text{-}1\ lock}$ for 90 minutes. *K. pneumoniae* 567880.1 was incubated in 10% C9-depleted serum supplemented with 80 nM $C9_{wt}$ or $C9_{TMH\text{-}1\ lock}$. a) OM damage was measured by $DiOC_2$ influx, which was determined by the shift in red:green fluorescence ratio over time in a multiwell plate-reader assay. b) IM damage was measured by Sytox influx over time in a multiwell plate-reader assay. Multiwell plate-reader assays are shown by one representative experiment that has been repeated at least three times.

**Supplementary figure 6 (S6)**

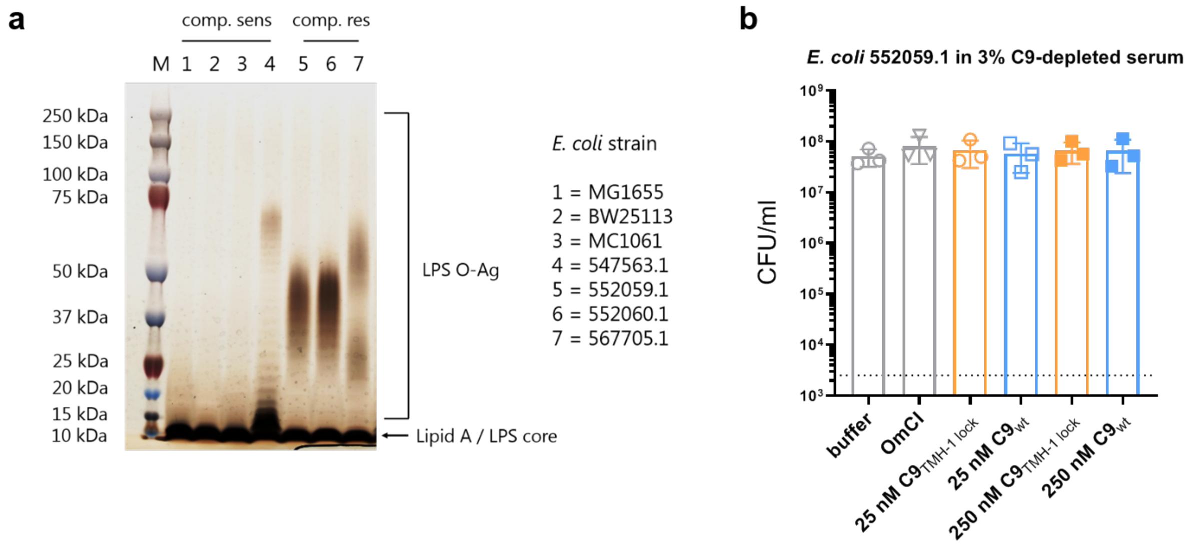


a) Complement-sensitive (MG1655, BW25113, MC1061 and 547563.1) and complement-resistant (552059.1, 552060.1, 567705.1) *E. coli* strains were typed for the presence of LPS O-Ag via Silver staining. Lipid A and LPS-core were distinguished from LPS O-Ag based on size. SDS-PAGE image is representative for at least two independent experiments. b) Complement-resistant *E. coli* 552059.1 was incubated in 3% normal human serum (NHS) supplemented with 25, 75 or 250 nM $C9_{wt}$, $C9_{TMH\text{-}1\ lock}$ or 25 µg/ml OmCI. Bacterial viability was determined by counting colony forming units (CFU’s) per ml. Data represent individual values with mean +/- SD of three independent experiments.
